# An Extended Primer Grip of Picornavirus Polymerase Facilitates Sexual RNA Replication Mechanisms

**DOI:** 10.1101/2019.12.13.876292

**Authors:** Brian J. Kempf, Colleen L. Watkins, Olve B. Peersen, David J Barton

## Abstract

Picornaviruses have both asexual and sexual RNA replication mechanisms. Asexual RNA replication mechanisms involve one parental template whereas sexual RNA replication mechanisms involve two or more parental templates. Because sexual RNA replication mechanisms counteract ribavirin-induced error catastrophe, we selected for ribavirin-resistant poliovirus to identify polymerase residues that facilitate sexual RNA replication mechanisms. We used serial passage in ribavirin, beginning with a variety of ribavirin-sensitive and ribavirin-resistant parental viruses. Ribavirin-sensitive virus contained an L420A polymerase mutation while ribavirin-resistant virus contained a G64S polymerase mutation. A G64 codon mutation (G64^Fix^) was used to inhibit emergence of G64S-mediated ribavirin resistance. Revertants (L420) or pseudo-revertants (L420V, L420I) were selected from all independent lineages of L420A, G64^Fix^ L420A and G64S L420A parental viruses. Ribavirin-resistant G64S mutations were selected in two independent lineages and novel ribavirin-resistance mutations were selected in the polymerase in other lineages (M299I, M323I, M392V, T353I). The structural orientation of M392, immediately adjacent to L420 and the polymerase primer grip region, led us to engineer additional polymerase mutations into poliovirus (M392A, M392L & M392V and K375R & R376K). L420A revertants and pseudorevertants (L420V, L420I) restored efficient sexual RNA replication mechanisms, confirming that ribavirin-induced error catastrophe coincides with defects in sexual RNA replication mechanisms. Viruses containing M392 mutations (M392A, M392L & M392V) and primer grip mutations (K375R & R376K) exhibited divergent RNA recombination, ribavirin sensitivity and biochemical phenotypes, consistent with changes in the fidelity of RNA synthesis. We conclude that an extended primer grip of the polymerase, including L420, M392, K375 & R376, contributes to the fidelity of RNA synthesis and to efficient sexual RNA replication mechanisms.

**IMPORTANCE:** Picornaviruses have both asexual and sexual RNA replication mechanisms. Sexual RNA replication shapes picornavirus species groups, contributes to the emergence of vaccine-derived polioviruses and counteracts error catastrophe. Can viruses distinguish between homologous and non-homologous partners during sexual RNA replication? We implicate an extended primer grip of the viral polymerase in sexual RNA replication mechanisms. By sensing RNA sequence complementarity near the active site, the extended primer grip of the polymerase has the potential to distinguish between homologous and non-homologous RNA templates during sexual RNA replication.

## INTRODUCTION

RNA viruses arose billions of years ago, becoming ubiquitous parasites of cellular life (1). The RNA-dependent RNA polymerase of RNA viruses is monophyletic, providing a means to consider the evolutionary history of all RNA viruses, to compare distinct groups of RNA viruses, and to classify RNA viruses (2, 3). Picornaviruses, whose origins predate the radiation of eukaryotic supergroups (4), have been studied extensively. Poliovirus, in particular, has been studied intensively for 100 years (5). The poliovirus RNA-dependent RNA polymerase was initially purified and biochemically characterized in 1977 (6). Since then, labs across the globe have studied the poliovirus RNA-dependent RNA polymerase in great detail [reviewed in (7)]. Picornavirus RNA-dependent RNA polymerases are multifunctional, catalyzing distinct steps of viral RNA replication, from making primers (VPg-uridylylation) (8–12), to replicating viral RNA (13, 14), to polyadenylation of progeny RNA genomes (15). Solving the atomic structure of poliovirus polymerase (16), and its elongation complex (17), led to unprecedented opportunities, including our ongoing work to understand viral RNA replication mechanisms and error catastrophe (18).

Picornaviruses have both asexual and sexual RNA replication mechanisms, both of which are catalyzed by the viral RNA-dependent RNA polymerase (Fig. 1). Asexual RNA replication mechanisms involve one parental template whereas sexual RNA replication mechanisms involve two or more parental templates. Previously, we established that an L420A polymerase mutation exacerbates ribavirin-induced error catastrophe coincident with defects in sexual RNA replication mechanisms (Fig. 1) (18, 19). Asexual RNA replication mechanisms are advantageous because vast amounts of progeny can be produced very quickly from one parental RNA template. However, asexual RNA replication mechanisms, in conjunction with error-prone RNA-dependent RNA polymerases (20, 21), can be disadvantageous, contributing to a loss of fitness due to Muller’s ratchet (22, 23). Errors introduced into viral RNA during asexual RNA replication cannot be easily removed in the absence of viral RNA recombination, except by reversion or negative selection (24). Reiterative asexual RNA replication, in conjunction with genetic bottlenecks, leads to error catastrophe, an overwhelming accumulation of mutations in viral RNA. Negative-strand RNA viruses like vesicular stomatitis virus, which do not recombine, are especially susceptible to Muller’s ratchet (23). In contrast, picornaviruses, which have relatively high levels of replicative RNA recombination (25, 26), are somewhat resistant to Muller’s ratchet (27–29). Nonetheless, ribavirin, a mutagenic antiviral drug, can restrict picornavirus replication via error catastrophe (30, 31). Replicative RNA recombination, a form of sexual RNA replication (32), can counteract ribavirin-induced error catastrophe (18, 19), presumably by purging lethal mutations from viral RNA genomes (33, 34).

**Figure 1.**
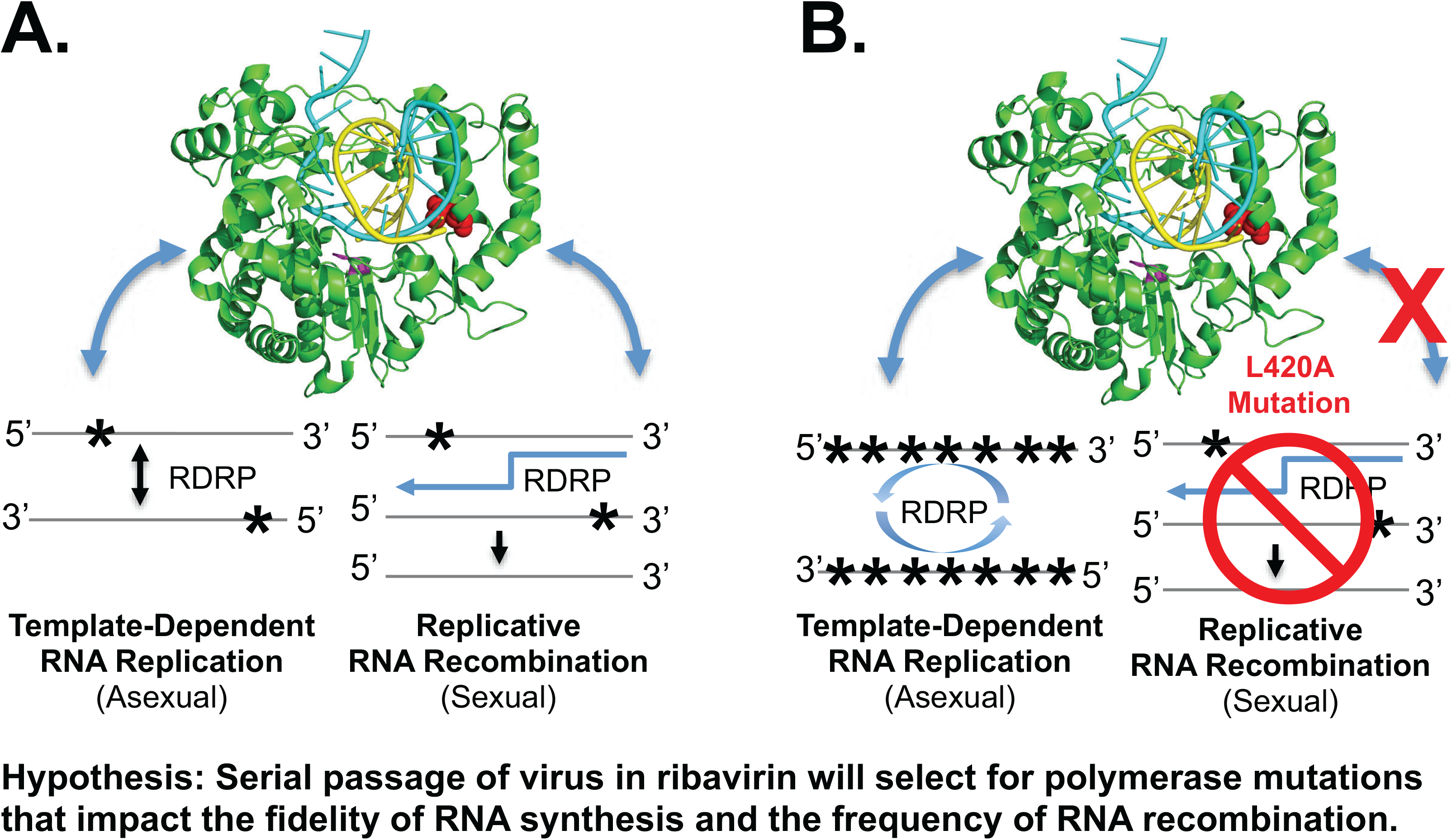
Picornaviruses have both asexual and sexual RNA replication mechanisms. (A) Asexual RNA replication involves one parental template whereas sexual RNA replication involves two parental templates. Mutations (asterisks) introduced during asexual RNA replication can be removed by sexual RNA replication. (B) Ribavirin-induced error catastrophe. The fidelity of RNA synthesis (53, 58) and the frequency of RNA recombination (18, 19) influence ribavirin-induced error catastrophe. Because an L420A mutation exacerbates ribavirin-induced error catastrophe coincident with defects in sexual RNA replication (18), we hypothesized that ribavirin-resistant poliovirus selected from L420A background would acquire novel mutations that restore efficient sexual RNA replication. Modified from Kempf et al. 2019 (18).

A growing body of evidence indicates that sexual RNA replication mechanisms play an essential role in picornavirus speciation and ongoing genetic exchange between viruses within defined species groups (35–41). Mucosal surfaces in the respiratory (42) and gastrointestinal tracts (41) are important ecological environments for ongoing genetic exchange between related picornaviruses. Gastrointestinal bacteria may even enhance virus co-infection to promote recombination (43). RNA sequence complementarity between genomes undergoing recombination (26, 44), among other factors (45), influence the frequency of recombination, with higher rates of recombination when sequences are more alike. When two related viruses co-infect a cell, sexual RNA replication mechanisms lead to the generation of chimeric RNA genomes, which are often fit, culminating in sustained host-to-host transmission. Circulating vaccine-derived polioviruses, obstacles to poliovirus eradication, are products of sexual RNA replication mechanisms – formed when unfit portions of OPV RNA genomes recombine with more fit regions of non-polio group C enteroviruses (46–48). When recombinant progeny are fit, as occurs in the case of circulating vaccine-derived polioviruses, the recombinant strains circulate from host-to-host (49, 50). Thus, an important biological consequence of sexual RNA replication mechanisms is the frequent exchange of genetic material among related picornaviruses. Picornavirus species groups are sustained over time by repeated, ongoing genetic exchange between related viruses within the species group. A leucine 420 residue in the polymerase thumb domain, conserved across picornavirus species groups, is required for efficient RNA recombination / sexual RNA replication (18, 19).

The interplay between asexual and sexual RNA replication (Fig. 1) is central to what Eigen deemed a grand challenge for the 21^st^ century, elucidating the complex mechanisms of error catastrophe, which are variable from one type of virus to another (51). In this study, we sought to identify features of the picornavirus polymerase involved in replicative RNA recombination / sexual RNA replication mechanisms. We reasoned that ribavirin-resistant virus might acquire polymerase mutations that facilitate sexual RNA replication mechanisms as a means to avoid ribavirin-induced error catastrophe. Building upon previous studies (18, 19), we used several ribavirin-sensitive and ribavirin-resistant parental viruses, in conjunction with serial passage in ribavirin, to select for ribavirin resistant poliovirus. Novel ribavirin resistance mutations, along with L420A revertants and pseudorevertants, were identified and characterized. These data implicate an extended primer grip of the viral polymerase in sexual RNA replication mechanisms.

## MATERIALS AND METHODS

### Poliovirus and infectious cDNA clones

Poliovirus type 1 (Mahoney), and mutant derivatives thereof, were derived from an infectious cDNA clone (52). Poliovirus RNAs were produced by T7 transcription of MluI-linearized cDNA clones (Ampliscribe T7, Cellscript Inc.) and transfected into HeLa cells to make infectious virus as previously described (15, 18, 19).

### Serial passage of poliovirus in escalating concentrations of ribavirin

Poliovirus was grown in HeLa cells in the presence of escalating concentrations of ribavirin (Sigma-Aldrich) using methods modified from those of Pfeiffer and Kirkegaard (53). HeLa cells were plated in 35mm 6-well dishes 24 hours before infection with WT or mutant parental strains of poliovirus at an MOI of 0.1 PFU per cell and incubated in 0 uM (P0), 100 uM (P1 to P4), 400 uM (P5 to P9) or 800 uM ribavirin (P10). Virus was harvested at 24 hpi by three freeze-thaw cycles and titered by plaque assay in HeLa cells. Three independent lineages of each parental strain were subjected to 10 serial passages in escalating concentrations of ribavirin. We expanded the population of poliovirus for each lineage after passage 10 by infecting a T150 flask of HeLa cells.

Viral cDNA was prepared from expanded passage 10 (P10^e^) viruses from each lineage and sequenced to identify polymerase mutations fixed in the virus populations.

### One-step growth of poliovirus

HeLa cells were plated in 35mm 6-well dishes 24 hours before infection with wildtype and mutant polioviruses. An MOI of 10 PFU per cell was used for one-step growth conditions. After 1 hour for virus adsorption, the inoculum was removed and the cells were incubated with 2 ml of culture media at 37°C. Total virus was harvested by three freeze-thaw cycles at the indicated times post-infection. Titers were determined by plaque assays. Mean titers from triplicates were plotted versus time post-infection with standard deviation error bars.

### Ribavirin dose-response assays

HeLa cells were plated in 35mm 6-well dishes 24 hours before infection with wildtype and mutant polioviruses. An MOI of 0.1 PFU per cell was used for ribavirin-dose response assays. After 1 hour for virus adsorption, the inoculum was removed and the cells were incubated with 2 ml of culture media at 37°C. Total virus was harvested by three freeze-thaw cycles at 24 hours post-infection. Titers were determined by plaque assays. Mean titers from triplicates were plotted with standard deviation error bars. Statistical significance was determined using the pairwise *t* test from GraphPad Prism (La Jolla, CA).

### Viral RNA recombination and replication controls

Viral RNA recombination assays and replication controls were performed as previously described (18, 19). For viral RNA recombination, L929 murine cells were co-transfected with two viral RNAs, each of which contained a lethal mutation: ΔCapsid Donor and 3D^pol^ ΔGDD Recipient. For replication controls, L929 cells were transfected with full-length infectious poliovirus RNAs containing wildtype or mutant polymerase, as indicated. Infectious poliovirus recovered from the transfected cells was titered by plaque assay in HeLa cells. Mean titers from triplicates were plotted with standard deviation error bars. Statistical significance was determined using the pairwise *t* test from GraphPad Prism (La Jolla, CA).

### Biochemical characterization of purified polymerase

Biochemical characteristics of purified polymerase were examined as previously described (54–57). Briefly, proteins were expressed in *E. coli* and purified through metal affinity, ion exchange, and gel filtration chromatography. Initiation rates are based on the time needed to form a +1 product after mixing 5 uM polymerase with 0.5 uM “10+1–12 RNA” and 40 uM GTP in reaction buffer containing 50 mM NaCl, 4 mM MgCl_2_, 25 mM HEPES (pH 6.5), and 2 mM Tris(2-carboxyethyl)-phosphine hydrochloride (TCEP), all at room temperature. Elongation complex stability measurements are based on diluting a 15-minute initiation reaction 10-fold into the same buffer with 300 mM NaCl and then testing the amount of elongation competent complex present at time points up to 4 hours and fitting the resulting data to a single exponential decay function. Kinetics assays were done using rapid stopped-flow fluorescent methods in 75 mM NaCl, 50 mM HEPES pH7, and MgCl_2_ in 4 mM excess over total NTP concentration. Processive elongation rates were determined with 26-nt long template RNA bearing a 5’-fluorescein end label (54) and the single cycle data were obtained with an RNA whose fluorescence reports on translocation of a template strand 2-aminopurine base from the +2 to +1 site following incorporation of a single CMP or 2’-dCMP (57). The Discrimination Factor is the ratio of catalytic efficiencies of CTP and dCTP incorporation reactions, i.e. (k_pol_/K_m_) CTP / (k_pol_/K_m_) dCTP.

## RESULTS

### Serial passage of poliovirus in ribavirin

We adapted the methods of Pfeiffer and Kirkegaard (53) to select for ribavirin resistance in poliovirus (Fig. 2). We used serial passage in escalating concentrations of ribavirin (Fig. 2A), beginning with several ribavirin-sensitive and ribavirin-resistant parental viruses (Fig. 2B): ribavirin-sensitive L420A virus, ribavirin-resistant G64S virus and virus with a G64 codon mutation (G64^Fix^) designed to inhibit emergence of G64S-mediated resistance. Altogether, this study contained eight parental strains with three independent lineages per strain: wildtype (WT), G64S, G64^Fix^, D79H, L419A, L420A, G64^Fix^ L420A and G64S L420A. Poliovirus with these mutations have normal one-step growth phenotypes in HeLa cells and all of the mutations are stably maintained in virus populations in the absence of ribavirin (18). A G64S mutation in the poliovirus polymerase mediates resistance to ribavirin by increasing the fidelity of RNA synthesis (53, 58) while an L420A mutation in the poliovirus polymerase increases sensitivity to ribavirin by inhibiting replicative RNA recombination / sexual RNA replication (18, 19). We used an MOI of 0.1 PFU per cell for each serial passage, harvesting virus by freeze thaw at 24 hpi. This MOI was high enough to avoid genetic bottlenecks; consequently, ribavirin-resistance mutations became fixed within the population only if they out-competed parental virus.

In order to maintain an MOI of 0.1 PFU per cell for each serial passage, we monitored the titer of poliovirus recovered from each infection (Fig. 3). P0 titers ~10^9^ PFU per ml were obtained in the absence of ribavirin (Fig. 3, P_0_). Under most circumstances, the titers for each lineage of virus were similar from passage to passage, with lower titers recovered as the concentrations of ribavirin increased: titers of 10^8^ to 10^9^ PFU per ml for passage 1-4 in 100 uM Rb, a wider range of titers from 10^5^ to 10^8^ PFU per ml for passage 5-9 in 400 uM Rb and titers ~ 10^6^ to 10^7^ PFU per ml for passage 10 at 800 uM Rb. In some cases, the titers of poliovirus recovered for independent lineages varied considerably from one passage to another: L420A lineage 2 (L420A2) titers dropped precipitously at passage 5 (~ 10^5^ PFU per ml) and remained low through passage 8 whereas L420A lineages 1 and 3 dropped at passage 5 but then increased incrementally from passage 5 through passage 9. Similar divergence was apparent in G64^Fix^ L420A lineages: G64^Fix^ L420A lineage 2 (G64^Fix^ L420A2) titers did not drop at passage 5 whereas G64^Fix^ L420A lineages 1 and 3 dropped precipitously at passage 5, with an incremental recovery by passage 9. In contrast, titers for G64S lineages 1-3 remained relatively high at all passages, as compared to other parental strains. Altogether, these data show that WT, ribavirin-resistant and ribavirin-sensitive parental strains behave differently during serial passage in escalating concentrations of ribavirin, and that individual lineages of ribavirin sensitive parental viruses become more resistant to ribavirin at different points during passage.

**Figure 2.**
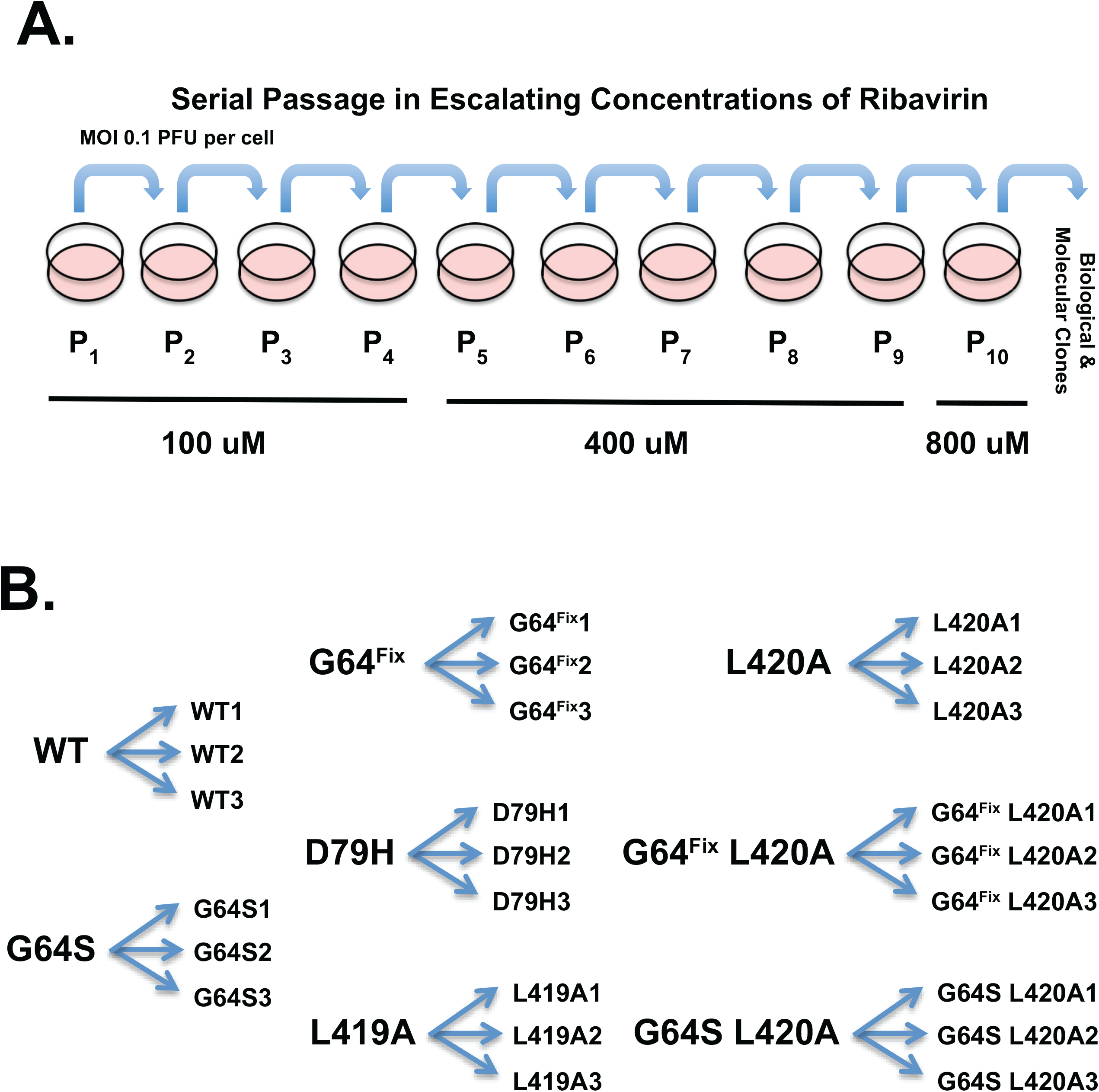
Selection of ribavirin-resistant poliovirus. (A) Serial passage of poliovirus in escalating doses of ribavirin. Methods adapted from Pfeiffer and Kirkegaard, 2003 (53). (B) Diagram showing the eight parental and 24 progeny virus strains used in this study.

**Figure 3.**
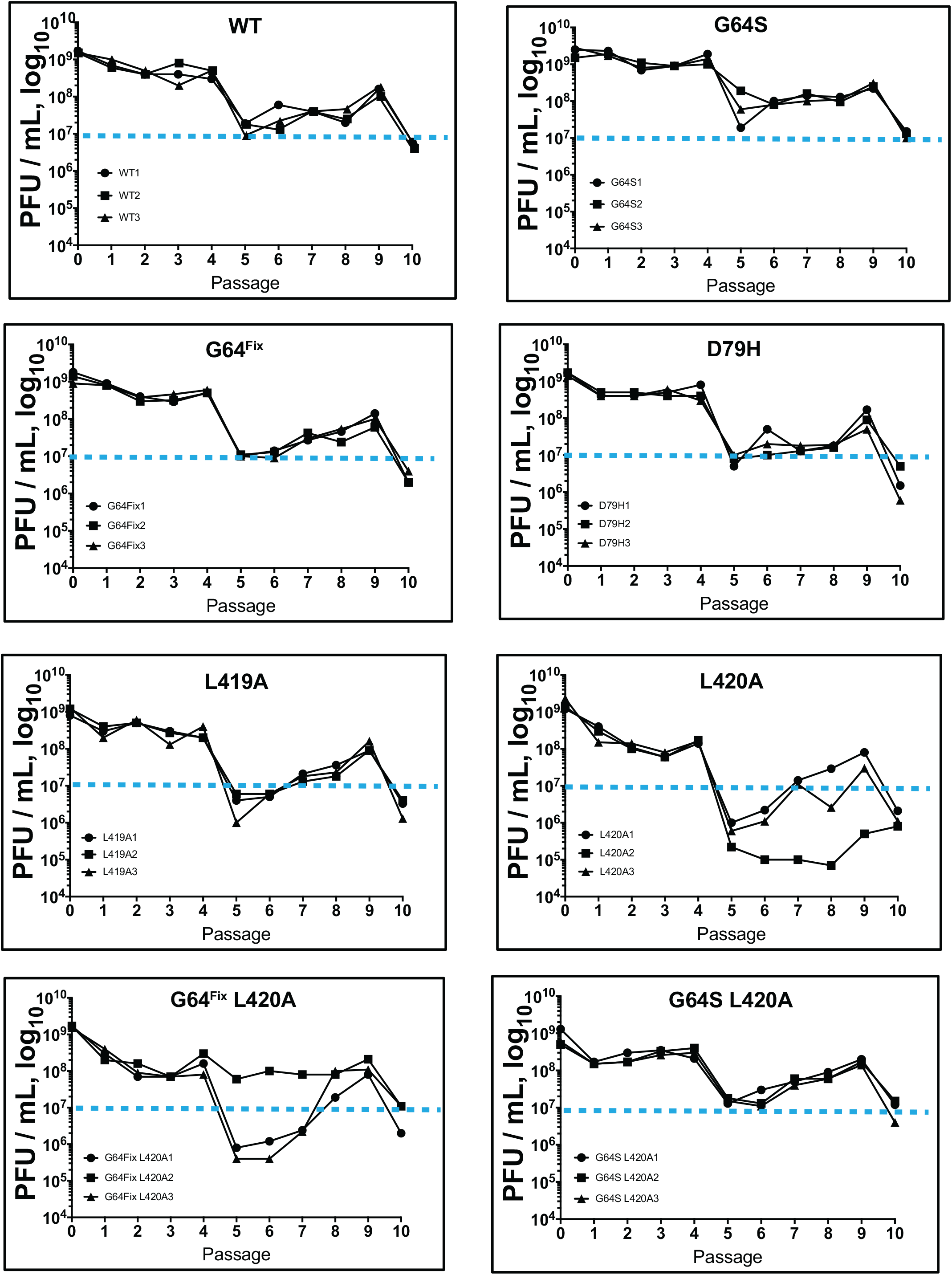
Titers of poliovirus recovered during serial passage in ribavirin. HeLa cells were infected with WT or mutant parental strains of poliovirus at an MOI of 0.1 PFU per cell, with three independent lineages per virus, and passaged 10 times (P1-P10). Virus was incubated in 0 uM (P0), 100 uM (P1 to P4), 400 uM (P5 to P9) or 800 uM ribavirin (P10). Virus was harvested at 24 hpi and titered by plaque assay at each passage. Dashed line provides reference point for 100-fold decrease in titers as compared to no drug at P_0_.

### Polymerase mutations selected by serial passage in ribavirin

We expanded the population of poliovirus for each lineage after passage 10 and sequenced viral cDNA to identify polymerase mutations fixed in the virus populations (Table 1). No polymerase mutations were selected in 12 of 24 lineages, including all lineages of WT and ribavirin-resistant G64S parental viruses. These data indicate that genetic bottlenecks were avoided - no mutations were detected in expanded passage 10 (P10^e^) populations from WT or G64S parental viruses. In contrast, we identified revertants, pseudorevertants or second site mutations fixed in expanded passage 10 (P10^e^) virus populations in 9/9 lineages of ribavirin-sensitive parental viruses (Table 1). Ribavirin-sensitive parental viruses include L420A, G64^Fix^ L420A and G64S L420A (18). Polymerase mutations were also detected in 1/3 lineages of G64^Fix^, D79H and L419A parental viruses, parental viruses with ribavirin sensitivity like WT virus (Table 1). The precise nature of polymerase mutations varied from one ribavirin-sensitive parental virus to another. A variety of polymerase mutations were selected from L420A and G64^Fix^ L420A parental viruses (Table 1. Lineage 1-3 of L420A and G64^Fix^ L420A viruses) while G64S L420A parental viruses had one singular outcome – reversion to L420 while maintaining the G64S resistance mutation (Table 1, G64S L420A Lineages 1-3). These data indicate that mutational robustness played a role in the nature of selected ribavirin-resistance mutations (34, 59). When G64S was present alone (G64S parental virus), or within an otherwise ribavirin-sensitive parental population (G64S L420A parental virus), the only outcome was L420A reversion to WT while maintaining G64S alleles. In contrast, when G64S was absent from the parental virus, or a codon mutation was used to inhibit G64S emergence, a variety of novel polymerase mutations were selected during serial passage in escalating concentrations of ribavirin. Thus, by using a variety of parental viruses, a variety of polymerase mutations were selected during serial passage in escalating concentrations of ribavirin. Altogether, polymerase mutations were selected in 12/24 lineages, including L420A revertants (L420), L420A pseudorevertants (L420V and L420I), G64S and several novel ribavirin resistance mutations (M299I, M323I, M392V and T353I) (Table 1).

**Table 1.**
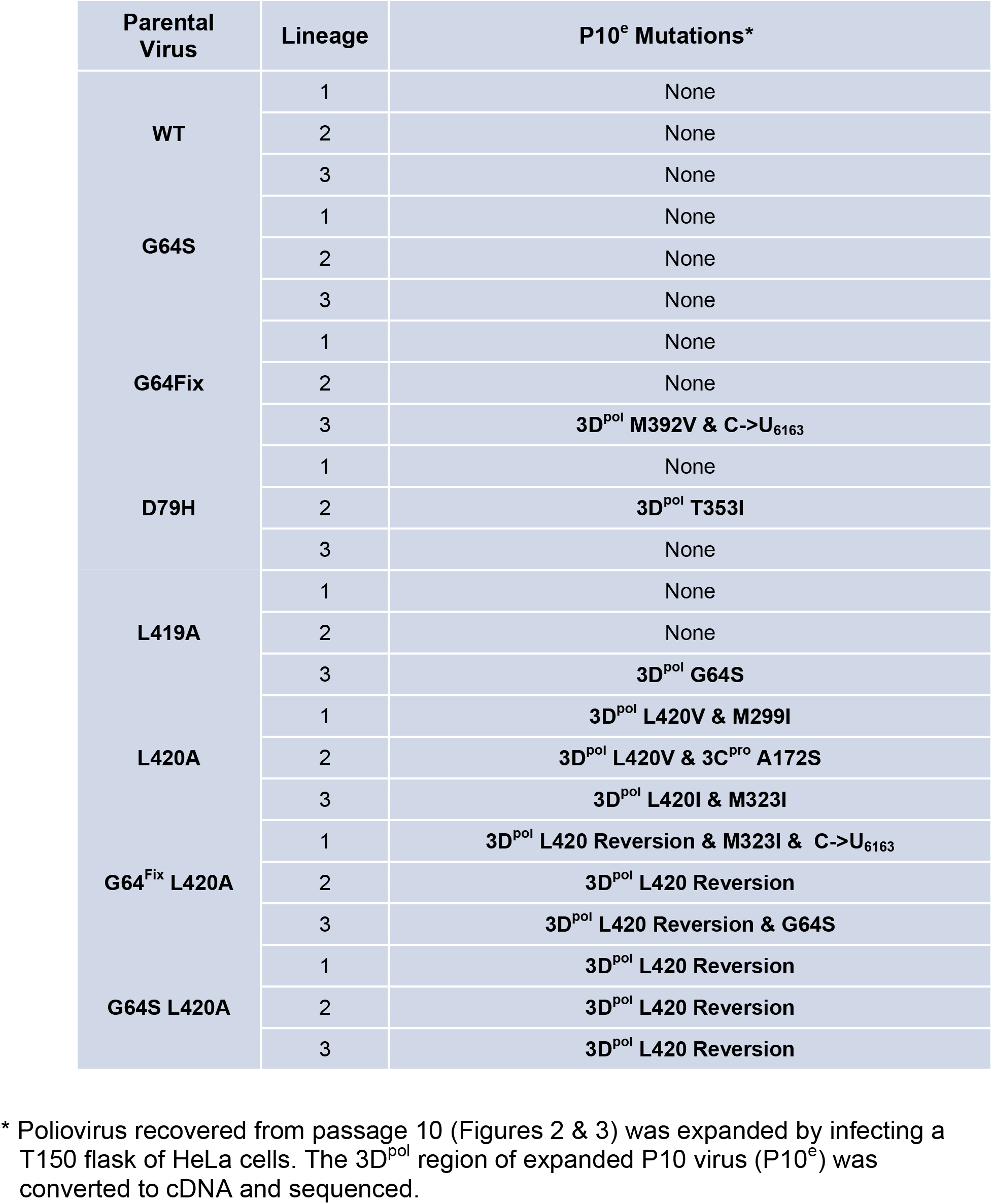
Polymerase mutations selected by serial passage in ribavirin.

The polymerase mutations selected during serial passage in escalating concentrations of ribavirin were located at several sites within polymerase elongation complexes, but all were in close proximity to RNA or the active site (Fig. 4). G64S, a well-studied ribavirin-resistance mutation (53, 58), is distal from the active site, near the back of the palm domain (Fig. 4. G64S). L420A revertants and pseudorevertants are present in an alpha-helix in the thumb region of the polymerase, where they pack into the minor groove of dsRNA products adjacent to the active site of the polymerase (Fig. 4, L420). L420, and presumably pseudorevertants thereof (L420V and L420I), interact with the ribose of viral RNA products 3 bases from the active site (19). M392V is found immediately adjacent to L420 and the primer grip in polymerase elongation complexes (Fig. 4, M392V). The location of the M392V ribavirin-resistance mutation underpins several important insights from this investigation. T353I is near the back of the palm domain, near G64 (Fig. 4, T353I). G64S and T353I residues both interact with S340, providing a common partner for transmitting allosteric effects from the back of the palm to the active site. M299I is buried in the palm domain, not far from M323I, which is found in the beta-sheet containing the active site (Fig. 4).

**Figure 4.**
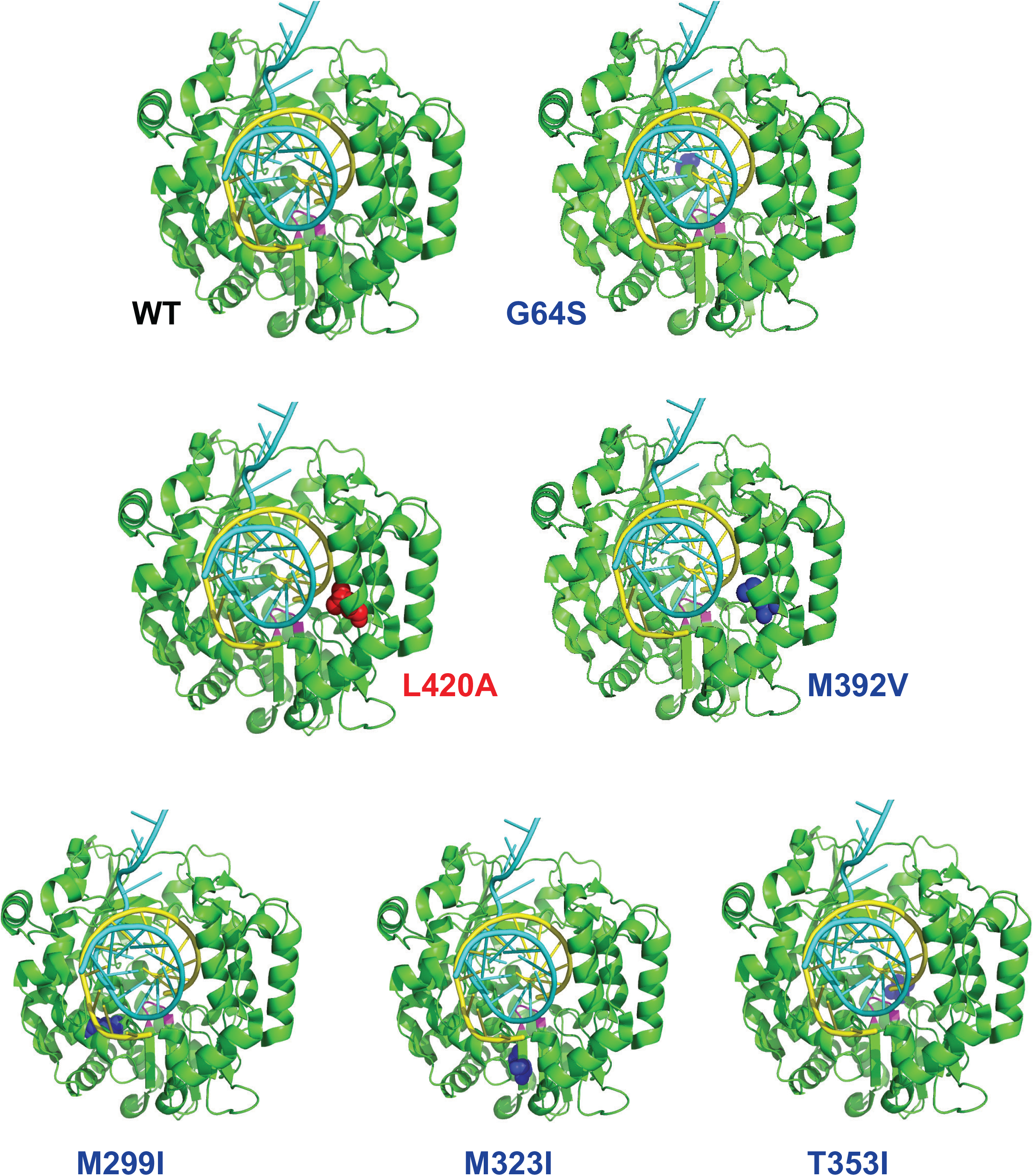
Location of polymerase mutations within elongation complexes. The atomic structure of poliovirus polymerase elongation complexes (17) was used to highlight the locations of mutations selected during serial passage in ribavirin. Poliovirus polymerase PDB entry 4K4T was rendered using PyMOL molecular graphics (Schrodinger, LLC). Polymerase is shown in green, RNA template in light blue, RNA products in yellow and the locations of amino acid changes during serial passage in ribavirin are highlighted in blue (G64S, M392V, T353I, M299I and M393I) or red (L420A) spheres.

### Ribavirin sensitivity of parental and progeny virus

We compared the titers of parental and progeny virus populations grown in the presence and absence of 600 uM ribavirin (Fig. 5). Expanded passage 10 (P10^e^) viruses from each lineage were compared to their respective parental viruses. Titers ~ 10^9^ PFU per ml were obtained for all viruses when they were grown in the absence of ribavirin (Fig. 5). WT poliovirus, and the P10^e^ viruses derived from WT after serial passage in escalating concentrations of ribavirin, remained sensitive to inhibition by ribavirin, with titers below 10^7^ PFU per ml when grown in 600 uM ribavirin (Fig. 5, WT). G64S parental virus, and the P10^e^ viruses derived therefrom, retained resistance to ribavirin, with titers well above 10^7^ PFU per ml when grown in the presence of 600 uM ribavirin (Fig. 5, G64S). Expanded passage 10 (P10^e^) virus from other parental strains exhibited a spectrum of sensitivity to ribavirin, as compared to WT and G64S viruses (Fig. 5). G64^Fix^ parental virus was similar to WT poliovirus; however, G64^Fix^ lineage 3 was more resistant to ribavirin (Fig. 5, G64^Fix^), presumably due to the selected M392V mutation (Table 1). D79H parental virus was similar to WT poliovirus, as reported (33), yet D79H lineage 2 was modestly resistant to ribavirin (Fig. 5, D79H panel), presumably due to the selected T353I mutation (Table 1). L419A parental virus was similar to WT poliovirus, as reported (18, 19), yet L419A lineage 3 was more resistant to ribavirin (Fig. 5, L419A panel), no doubt due to the selected G64S mutation (Table 1). L420A parental virus was more sensitive to ribavirin than WT virus, as reported (18, 19); however, L420A lineage 1, 2 and 3 viruses grew to higher titers in the presence of ribavirin (Fig. 5, L420A panel), presumably due to the selected polymerase mutations in each lineage (Table 1). G64^Fix^ L420A parental virus was more sensitive to ribavirin than WT virus, as reported (18, 19); however, G64^Fix^ L420A lineage 1, 2 and 3 viruses grew to higher titers in the presence of ribavirin (Fig. 5, G64^Fix^ L420A panel), presumably due to the selected polymerase mutations in each lineage (Table 1). Similarly, ribavirin-sensitive G64S L420A parental virus and its selected progeny exhibited ribavirin phenotypes consistent with the genotypes of the respective virus populations (Fig. 5 and Table 1). Altogether, these data suggest that the polymerase mutations selected during serial passage in escalating concentrations of ribavirin (Table 1) correlate with measurable resistance to ribavirin (Fig. 5).

**Figure 5.**
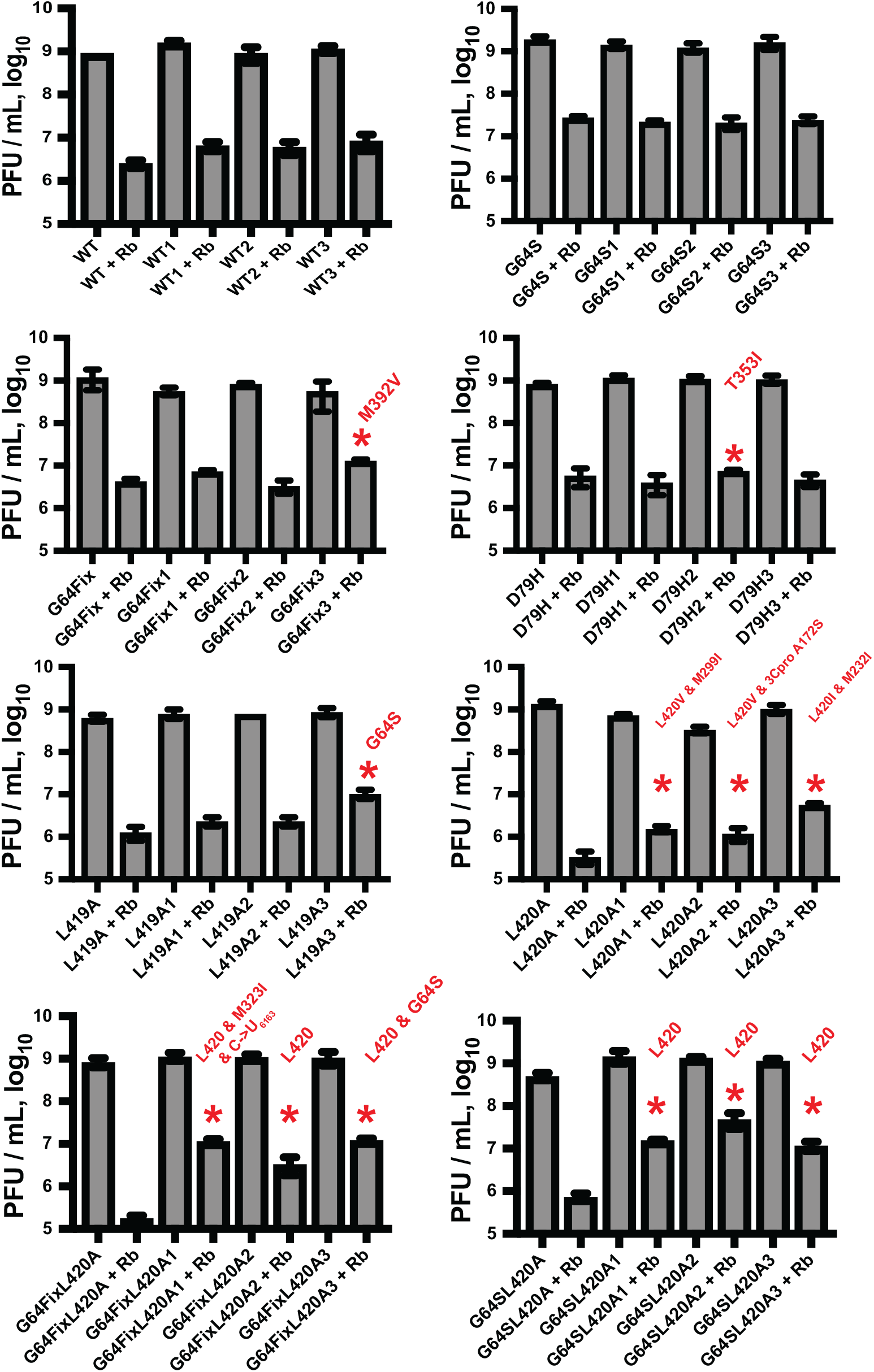
Ribavirin sensitivity of parental and progeny virus. Titer of parental and progeny virus grown in the absence and presence of 600 uM ribavirin. Expanded passage 10 (P10^e^) viruses from each lineage were used to infect HeLa cells at an MOI of 0.1 PFU per cell. Infected cells were incubated for 24 hrs in the absence or presence of 600 uM ribavirin. Virus was harvested by freeze-thaw and titered by plaque assay on HeLa cells. The names of parental strains and lineage 1, 2 and 3 strains are indicated on the X-axis. Polymerase mutations selected during serial passage and identified in Table 1 are annotated on the graphs accordingly (in red).

### Virus derived from infectious cDNA clones

The polymerase mutations identified during serial passage in escalating concentrations of ribavirin were reverse engineered into infectious cDNA clones of poliovirus to derive genetically defined virus populations (Table 2, Ribavirin Selected Mutations). In addition, several engineered variants originating from our work, and that of others, were similarly generated (Table 2, Engineered Variants) (18, 19, 53, 60). Many of these mutations lie within an extended primer grip region of the polymerase, consisting of L420, M392 and primer grip residues (K375 & R376). All of the mutations listed here were stably maintained in infectious virus, except for alanine substitution mutations in the primer grip residues (K375A & R376A). K375A and R376A mutations were unstable, reverting back to wildtype residues, precluding further evaluation, but the charge retaining K375R and R376K mutants at these sites were stable. Virus containing polymerase mutations were assessed in one-step growth assays (Fig. 6), ribavirin dose-response assays (Fig. 7A), viral RNA recombination assays (Fig. 7B) and replication controls (Fig. 7C).

**Figure 6.**
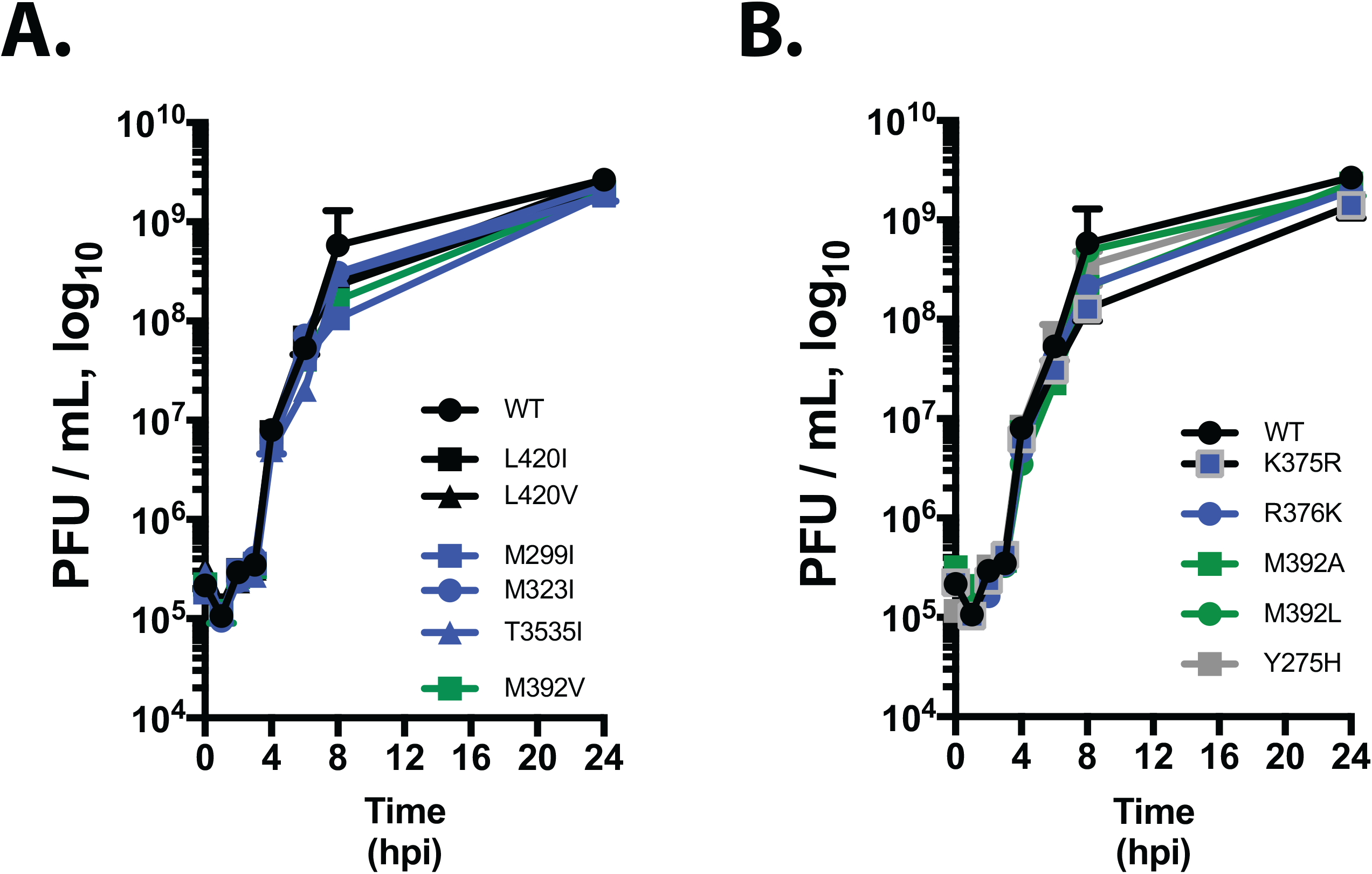
One-step growth of wildtype and mutant polioviruses. Polymerase mutations selected by serial passage in ribavirin were engineered into infectious cDNA clones. Viruses derived from infectious cDNA clones were compared under one-step growth conditions: HeLa cells were infected with wild-type or mutant poliovirus at an MOI of 10 PFU per cell. Virus was harvested at the indicated times by freeze-thawing cells. Titers were determined by plaque assay and plotted versus time (hpi, hours post-infection).

**Figure 7.**
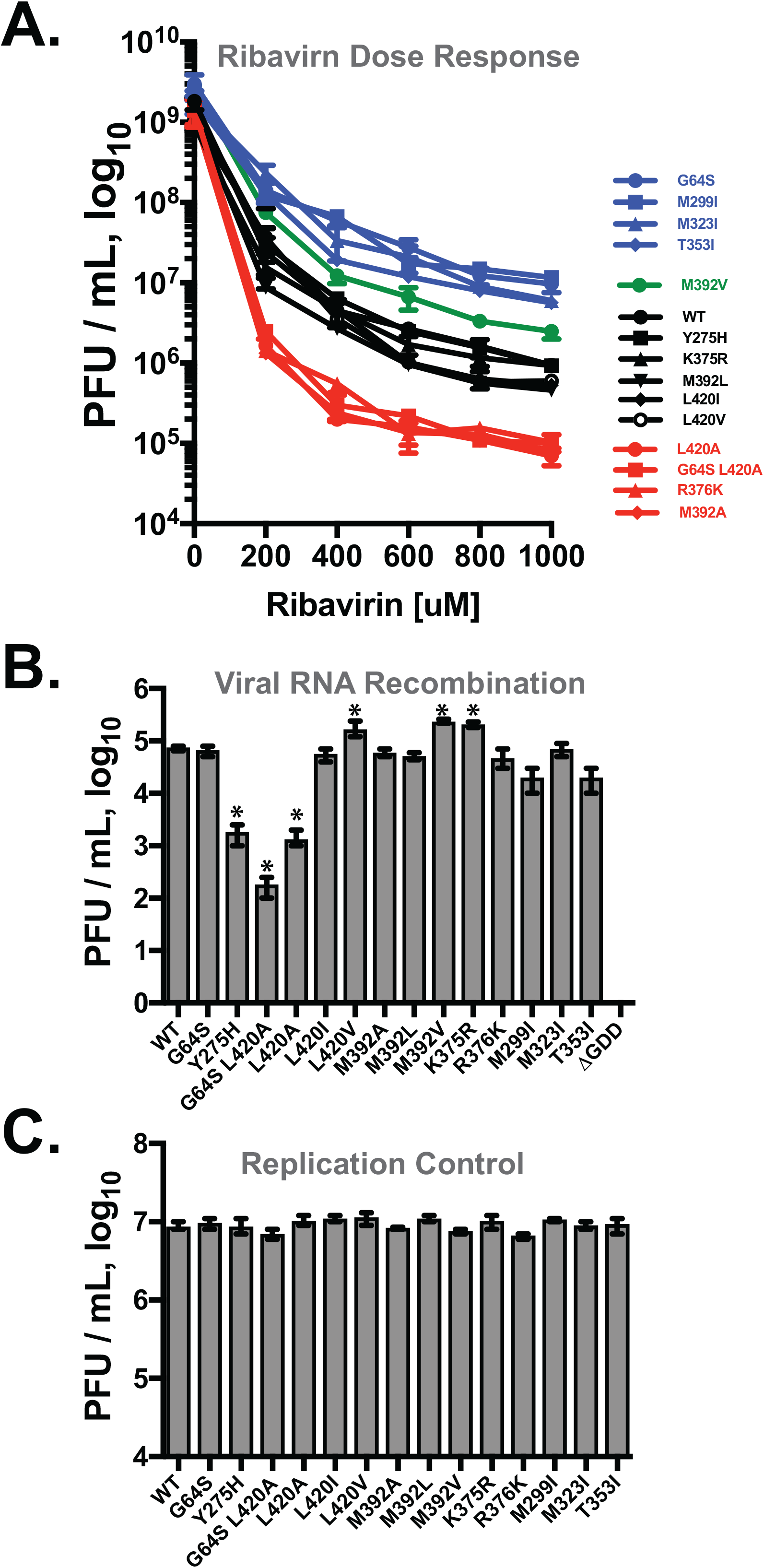
Polymerase mutations influence ribavirin-induced error catastrophe and sexual RNA replication mechanisms. (A) Polymerase mutations influence ribavirin sensitivity and resistance. HeLa cells were infected with wild-type or mutant poliovirus at an MOI of 0.1 PFU per cell and incubated for 24 h with the indicated concentrations of ribavirin. Each condition was performed in triplicate. After three freeze-thaw cycles, viral titers were determined by plaque assay and plotted versus ribavirin concentration. Four ribavirin-responsive phenotypic clusters were observed: a ribavirin-resistant cluster (blue), a modestly ribavirin-resistant virus (green), a wildtype cluster (black) and a ribavirin-sensitive cluster (red). *P* values are <0.006 between all phenotypic clusters. (B) Polymerase mutations influence the frequency of viral RNA recombination mechanisms, i.e. sexual RNA replication. ΔCapsid “Donor” and ΔGDD “Recipient” RNAs were co-transfected into murine cells as previously established (18, 19). ΔCapsid Donor contained wild-type or mutant polymerases as indicated on the x axis. The titer of virus produced in co-transfected murine cells was determined by plaque assays in HeLa cells. *, *P* values are <0.05 compared to the wildtype. (C) Replication controls showing titers from poliovirus RNA containing polymerase mutations (x-axis) after transfection into murine cells. At 72 hpt, the amount of poliovirus produced within the murine cells was determined by plaque assay.

**Table 2.** Virus derived from infectious cDNA.

| Polymerase* | Ribavirin Selected | Engineered Variant | Stably Maintained | Origin |
| --- | --- | --- | --- | --- |
| WT |  |  | Yes | Collis et al. 1992 |
| G64S |  |  | Yes | Pfeiffer and Kirkegaard, 2003 |
| L420A |  |  | Yes | Kempf et al. 2016 |
| G64S L420A |  |  | Yes | Kempf et al. 2019 |
| L420V |  |  | Yes | This study |
| L420I |  |  | Yes | This study |
| M392A |  |  | Yes | This study |
| M392L |  |  | Yes | This study |
| M392V |  |  | Yes | This study |
| K375R |  |  | Yes | This study |
| R376K |  |  | Yes | This study |
| K375A |  |  | No | This study |
| R376A |  |  | No | This study |
| M299I |  |  | Yes | This study |
| M323I |  |  | Yes | This study |
| T353I |  |  | Yes | This study |
| Y275H |  |  | Yes | Acevedo et al. 2018 |

\* Polymerase mutations identified during ribavirin selection, along with engineered variants, were cloned into infectious cDNA. Poliovirus was recovered from HeLa cells transfected with RNA derived from each clone. cDNA from the 3D<sup>pol</sup> region was sequenced to determine whether mutations were stably maintained in virus. Citations (18, 19, 52, 53, 60).

### One-step growth of wildtype and mutant polioviruses

Before assessing ribavirin sensitivity and resistance, we measured one-step growth of wildtype and mutant polioviruses in HeLa cells (Fig. 6). HeLa cells were infected at an MOI of 10 PFU per cell, and virus yields were determined at 0, 1, 2, 3, 4, 6, 8 and 24 hpi (Fig. 6). Each of the viruses grew by 4 orders of magnitude between 3 and 24 hpi, reaching titers ~10^9^ PFU per ml. These data indicate that the wildtype and mutant viruses have similar one-step growth phenotypes.

### Ribavirin sensitivity and resistance

Poliovirus-infected HeLa cells were used to assess ribavirin sensitivity and resistance (Fig. 7A). Wildtype and mutant polioviruses reached titers ~10^9^ PFU per ml in the absence of ribavirin (Fig. 7A, 0 uM ribavirin). Wildtype poliovirus titers decreased incrementally as the dose of ribavirin increased, reaching titers ~ 2 x 10^6^ PFU per ml in 1000 uM ribavirin (Fig. 7A, WT). Poliovirus with a G64S polymerase mutation resisted inhibition by ribavirin, as previously reported (18, 19, 53), reaching titers ~ 2 x 10^7^ PFU per ml in 1000 uM ribavirin (Fig. 7A, G64S). In contrast, poliovirus with an L420A polymerase mutation was more sensitive to inhibition by ribavirin, reaching titers ~ 1 x 10^5^ PFU per ml in 1000 uM ribavirin (Fig. 7A, L420A) (18, 19). Likewise, poliovirus containing both G64S and L420A mutations was more sensitive to ribavirin, reaching titers ~ 1 x 10^5^ PFU per ml in 1000 uM ribavirin (Fig. 7A, G64S L420A). Thus, the ribavirin-resistant (G64S) and ribavirin-sensitive (L420A) controls behaved as previously reported (18, 19).

The viruses used in this investigation segregated into four ribavirin-responsive phenotypic groups (Fig. 7A): a ribavirin-resistant cluster (blue), a modestly ribavirin-resistant virus (green), a wildtype cluster (black) and a ribavirin-sensitive cluster (red). Ribavirin-resistant polioviruses included those with G64S, M299I, M232I and T353I (Fig. 7A, ribavirin-resistant cluster in blue). Poliovirus with an M392V mutation was modestly resistant to ribavirin, as compared to the more ribavirin-resistant G64S cluster. Ribavirin-sensitive polioviruses included those with L420A, G64S L420A, R376K and M392A polymerase mutations. Polioviruses in the wildtype cluster included WT, Y275H, K375R, M392L, L420I and L420V. These results indicate that the polymerase mutations selected during serial passage in escalating concentrations of ribavirin (G64S, L420V, L420I, M392V, M299I, M323I, and T353I) are indeed ribavirin-resistance mutations, as compared to their parental strains. While the selected M392V mutation rendered poliovirus modestly resistant to ribavirin, an M392A variant was ribavirin-sensitive and an M392L variant had wt sensitivity to ribavirin. L420A and G64S L420A parental strains were ribavirin-sensitive viruses, whereas L420V and L420I containing viruses exhibited wildtype sensitivity to ribavirin. Conservative substitutions in the primer grip exhibited divergent ribavirin phenotypes: poliovirus containing a K375R mutation had wildtype sensitivity to ribavirin whereas poliovirus containing a R376K mutation was ribavirin-sensitive. These data indicate that polymerase mutations influence ribavirin sensitivity and resistance, with some mutations rendering virus more resistant to ribavirin while others render virus more sensitive to ribavirin.

### Viral RNA recombination

Based on our previous investigation (18), we predicted that ribavirin-resistant poliovirus might acquire polymerase mutations that facilitate sexual RNA replication mechanisms as a means to avoid ribavirin-induced error catastrophe. Consequently, we assayed the impact of polymerase mutations on the frequency of replicative RNA recombination (Fig. 7B). In replicative RNA recombination assays, we co-transfect murine cells with two viral RNAs, each of which contains a lethal mutation: 1, a subgenomic replicon with an in-frame capsid deletion carrying the mutant polymerase, and 2, a full-length poliovirus RNA with a lethal deletion of the active site GDD motif in the polymerase (18, 19). Wildtype or mutant polymerase is produced from the subgenomic replicon in the co-transfected cells. The amount of infectious poliovirus produced in the co-transfected murine cells corresponds with the frequency of viral RNA recombination. By comparing the titers of poliovirus, we assess the impact of polymerase mutations on the frequency of viral RNA recombination (Fig. 7B).

WT polymerase established wildtype levels of replicative RNA recombination in our experiments, with titers of poliovirus ~ 10^5^ PFU per ml (Fig. 7B, WT). A G64S mutation had no impact on the frequency of viral RNA recombination (Fig. 7B, G64S), as compared to WT polymerase. In contrast, an L420A mutation inhibited replicative recombination by two orders of magnitude, with poliovirus titers ~10^3^ PFU per ml (Fig. 7B, L420A). Likewise, a G64S L420A polymerase supported decreased levels of replicative RNA recombination, with titers near 10^2^ PFU per ml (Fig. 7B, G64S L420A). These outcomes for WT, G64S and L420A polymerases are similar to those previous reported (18): WT and G64S polymerases support wildtype magnitudes of viral RNA recombination whereas an L420A polymerase mutation significantly inhibits replicative RNA recombination.

L420A pseudorevertants (L420V and L420I), selected during serial passage in escalating concentrations of ribavirin, restored replicative RNA recombination frequencies to wildtype levels (Fig. 7B, L420I and L420V). All the other polymerase mutations selected during serial passage in escalating concentrations of ribavirin maintained replicative RNA recombination frequencies at near wildtype levels, albeit with some variation in titers both above and below 10^5^ PFU per ml (Fig. 7B, M392V, M299I, M323I, M392V and T353I).

Engineered polymerase mutations had divergent ribavirin and recombination phenotypes: M392A was ribavirin-sensitive with wildtype levels of recombination while M392L exhibited wildtype ribavirin sensitivity and wildtype levels of recombination (Fig. 7, M392A and M392L). Likewise, K375R exhibited wildtype ribavirin sensitivity and wildtype levels of recombination whereas R376K was ribavirin sensitive with wildtype levels of recombination (Fig. 7, K375R and R376K). Finally, consistent with data reported by Acevedo et al. (60), a Y275H mutation inhibited viral RNA recombination without impacting ribavirin sensitivity or resistance (Fig. 7A and 7B).

As a control for the viral RNA recombination assay, we examined the impact of polymerase mutations on virus replication in murine cells (Fig. 7C, Replication Controls). Full-length infectious RNA containing each of the polymerase mutations (Fig. 7C, x-axis) was transfected into murine cells. Infectious poliovirus was recovered at 72 hpi by freeze-thaw and titered by plaque assay (Fig. 7C, Replication Control). These data show that the polymerase mutations do not inhibit virus replication within murine cells. Consequently, the decreased titers of virus recovered from viral RNA recombination assays of Y275H, L420A and G64S L420A polymerases are due to defects in sexual RNA replication mechanisms (i.e. viral RNA recombination) rather than defects in asexual RNA replication mechanisms.

Altogether, these results indicate that ribavirin resistant poliovirus selected during serial passage in escalating concentrations of ribavirin restored (or maintained) efficient viral RNA recombination mechanisms. Furthermore, polymerase with engineered amino acid substitution mutations at M392 and primer grip residues (K375 and R376) exhibited divergent ribavirin and recombination phenotypes, with some mutations increasing sensitivity to ribavirin without negatively impacting viral RNA recombination frequencies.

### Biochemical phenotypes of wildtype and mutant polymerase

Having examined viral RNA recombination and ribavirin sensitivity, we next examined the biochemical phenotypes of wildtype and mutant polymerases (Table 3). We used purified polymerase to assess the effects of individual mutations on various biochemical parameters, including RNA synthesis initiation, elongation complex stability, processive elongation rate, and single nucleotide addition cycle rate. The fidelity of nucleotide addition was indirectly assayed via a nucleotide Discrimination Factor that is based on the relative catalytic efficiency of rNTP versus 2’-dNTP incorporation (61).

**Table 3.** Biochemical phenotypes of purified polymerase.

| Polymerase* | Initiation (min) | EC Stability (Min) | Processive Rate (nt/sec) | Processive $K_m$ (μM) | Single NTP Rate (per sec) | Single NTP $K_m$ (μM) | Discrimination Factor |
| --- | --- | --- | --- | --- | --- | --- | --- |
| From Kempf et al. 2019** |  |  |  |  |  |  |  |
| WT | 6 ± 1 | 88 ± 9 | 22 ± 1 | 67 ± 2 | 30 ± 1 | 37 ± 2 | 120 ± 10 |
| G64S | 8 ± 1 | 24 ± 7 | 12 ± 1 | 46 ± 4 | 23 ± 1 | 39 ± 5 | 220 ± 40 |
| L420A | 8 ± 1 | 28 ± 4 | 30 ± 1 | 75 ± 3 | 40 ± 1 | 39 ± 3 | 100 ± 10 |
| G64S L420A | 6 ± 3 | 40 ± 8 | 26 ± 1 | 84 ± 9 | 28 ± 1 | 36 ± 6 | 220 ± 50 |
| From this study |  |  |  |  |  |  |  |
| WT | 4 ± 1 | 130 ± 20 | 26 ± 1 | 72 ± 4 | 40 ± 1 | 46 ± 3 | 100 ± 10 |
| L420V | 4 ± 1 | 170 ± 10 | 29 ± 1 | 82 ± 5 | 41 ± 1 | 40 ± 3 | 99 ± 9 |
| L420I | 4 ± 1 | 60 ± 2 | 33 ± 1 | 90 ± 5 | 45 ± 1 | 41 ± 2 | 107 ± 8 |
| M392A | 3 ± 1 | 20 ± 1 | 30 ± 1 | 90 ± 8 | 33 ± 1 | 38 ± 3 | 120 ± 10 |
| M392L | 4 ± 1 | 39 ± 4 | 25 ± 1 | 74 ± 9 | 38 ± 1 | 34 ± 1 | 102 ± 6 |
| M392V | 4 ± 1 | 11 ± 1 | 22 ± 1 | 74 ± 7 | 42 ± 1 | 51 ± 3 | 160 ± 10 |
| K375R | 5 ± 1 | 51 ± 2 | 20 ± 1 | 96 ± 8 | 36 ± 1 | 44 ± 3 | 200 ± 20 |
| R376K | 2 ± 1 | 30 ± 3 | 65 ± 2 | 160 ± 10 | 48 ± 1 | 46 ± 3 | 17 ± 1 |
| M299I | 4 ± 1 | 180 ± 30 | 18 ± 1 | 66 ± 4 | 35 ± 1 | 42 ± 2 | 120 ± 10 |
| M323I | 4 ± 1 | 150 ± 20 | 18 ± 1 | 64 ± 4 | 37 ± 1 | 50 ± 3 | 130 ± 10 |
| T353I | 5 ± 1 | 90 ± 20 | 18 ± 1 | 73 ± 2 | 38 ± 1 | 49 ± 3 | 160 ± 10 |
| Y275H | 13 ± 6 | 95 ± 3 | 27 ± 1 | 95 ± 8 | 35 ± 1 | 42 ± 2 | 140 ± 10 |

\* Polymerase was expressed and purified. Biochemical phenotypes were determined as previously reported (55, 56).
\*\*Kempf et al. (2019) (18), with permission.

In a previous study (18), we compared the biochemical phenotypes of WT, G64S, L420A and G64S L420A polymerases (Table 3). A G64S mutation reduced the rate of RNA elongation in both single nucleotide (23 per second for G64S versus 30 per second for WT) and processive elongation assays (12 nt/s for G64S versus 22 nt/s for WT), with a corresponding increase in the fidelity of RNA synthesis (Discrimination Factor of 220 ± 40 for G64S polymerase versus 120 ± 10 for WT). Consequently, G64S is considered a high-fidelity polymerase (53, 58). An L420A mutation decreased elongation complex stability, increased RNA elongation rates, and exhibited a wildtype discrimination factor (Discrimination Factor of 100 ± 10 for L420A polymerase). Thus, L420A polymerase is considered to have normal (wildtype) fidelity. Polymerase containing both G64S and L420A mutations has hybrid phenotypes, with faster elongation rates like L420A polymerase and an increased discrimination factor (Discrimination Factor of 220 ± 50 for G64S L420A polymerase). Consequently, G64S L420A is considered a high-fidelity polymerase, like G64S polymerase. Consistent with our previous report (18), an L420A mutation inhibits viral RNA recombination (Fig. 7B, L420A) and renders virus more susceptible to ribavirin-induced error catastrophe (Fig. 7A, L420). Furthermore, a high-fidelity G64S L420A polymerase fails to render virus resistant to ribavirin when viral RNA recombination is disabled (Fig. 7, G64S L420A). Thus, a high fidelity polymerase fails to mediate ribavirin resistance when viral RNA recombination is disabled (Fig. 7, G64S L420A). These observations for WT, G64S, L420A and G64S L420A polymerases provide a foundational framework to interpret the biochemical data for the new polymerase mutants from this study (Table 3).

As compared to WT polymerase, L420V, M299I and T353I polymerases form more stable elongation complexes (Table 3, L420V, M299I and T353I). In contrast, mutations in the extended primer grip of the polymerase significantly reduced elongation complex stability (Table 3, L420A, L420I, M392A, M392L, M392V, K375R and R376K). The M392V mutation reduced elongation complex stability by 10-fold (EC Stability of 11 ± 1 min for M392V versus 130 ± 20 min for WT).

Primer grip mutations (K375R & R376K) exhibited divergent biochemical phenotypes (Table 3). The K375R mutation reduced the rate of RNA elongation in both single nucleotide (36 per sec for K375R versus 40 per sec for WT) and processive elongation assays (20 nt per sec for K375R versus 26 nt per sec for WT), with a corresponding increase in the fidelity of RNA synthesis (Discrimination Factor of 200 ± 20 for K375R polymerase). In contrast, the R376K mutation increased the rate of RNA elongation in both single nucleotide (48 per sec for R376K versus 40 per sec for WT) and processive elongation assays (65 nt per sec for R376K versus 26 nt per sec for WT), with a corresponding decrease in the fidelity of RNA synthesis (Discrimination Factor of 17 ± 1 for R376K polymerase). The low fidelity R376K polymerase rendered poliovirus more susceptible to ribavirin (Fig. 7A, R376K), as one might expect, however, the high fidelity K375R polymerase exhibited normal (wildtype) sensitivity to ribavirin (Fig. 7A, K375R). The R376K polymerase also initiated RNA synthesis 2-fold faster than WT polymerase. Together, these data show that conservative substitution mutations in the primer grip (K375R & R376K) result in divergent biochemical phenotypes (Table 3) and divergent ribavirin sensitivity (Fig. 7A).

Polymerase mutations selected during serial passage in ribavirin tend to have reduced rates of RNA elongation and increased discrimination factors (Table 3, G64S, M392V, M299I, M323I and T353I). The M299I, M323I and T353I mutations reduced the rate of RNA elongation in processive elongation assays (18 nt per sec for M299I, M323I and T353I versus 26 nt per sec for WT), with corresponding increases in the fidelity of RNA synthesis (Discrimination Factors of 120 ± 10 for M299I polymerase, 130 ± 10 for M323I polymerase and 160 ± 10 for T353I polymerase). The M392V mutation reduced the rate of RNA elongation in processive elongation assays (22 nt per sec for M392V versus 26 nt per sec for WT) and increased the fidelity of RNA synthesis (Discrimination Factor of 160 ± 10 for M392V polymerase). Increases in the fidelity of RNA synthesis are consistent with increased resistance to ribavirin (Fig. 7A, G64S, M392V, M299I, M323I and T353I). Altogether, these data indicate that the polymerase mutations selected during serial passage in escalating concentrations of ribavirin tend to increase the fidelity of RNA synthesis coincident with decreased rates of RNA elongation.

We included a Y275H mutation in this study based on the report from Acevedo et al. (2018) (60). Consistent with their data, we find that the Y275H mutation inhibits viral RNA recombination (Fig. 7B) without increasing sensitivity to ribavirin (Fig. 7A). These phenotypes are distinct from those associated with an L420A polymerase mutation, where decreased viral RNA recombination is mechanistically linked with increased sensitivity to ribavirin (Fig. 7A and 7B, L420A). The biochemical phenotypes of Y275H are important in this regard (Table 3). The Y275H mutation inhibits the initiation of RNA synthesis by 3-fold (13 ± 6 min for Y275H versus 4 ± 1 min for WT). None of the other mutations in our panel exhibit this phenotype. The proximity of Y275 to the template entry channel of the polymerase (56) suggests this defect in the initiation of RNA synthesis may arise from defects in template RNA binding. In the discussion, we elaborate on the structural and functional distinctions of Y275 and L420 residues in the polymerase, especially as they relate to viral RNA recombination and ribavirin-induced error catastrophe.

## DISCUSSION

In this investigation, we found that serial passage of poliovirus in escalating concentrations of ribavirin selected for ribavirin-resistant polymerase mutations that facilitate efficient replicative RNA recombination (Fig. 7) and increased fidelity of RNA synthesis (Table 3). These data reinforce and build upon prior studies regarding ribavirin-induced error catastrophe (18, 30, 51, 53). By using several ribavirin-sensitive and ribavirin-resistant parental viruses, serial passage in escalating doses of ribavirin led to the selection of both known and unknown ribavirin resistance mutations (Table 1). L420A revertants and pseudorevertants (L420I & L420V) regained efficient viral RNA recombination coincident with ribavirin resistance (Fig. 7). These data reinforce our previous conclusions that viral RNA recombination counteracts error catastrophe (18). A G64S mutation was selected in two instances, reinforcing its well-established role as a modulator of RNA synthesis fidelity (53, 58). In addition to G64S, four novel ribavirin-resistance mutations were identified: M299I, M323I, T353I and M392V. Two important phenotypes were shared by all of these ribavirin-resistant polymerase mutations: efficient viral RNA recombination and increased fidelity of RNA synthesis. Thus, both asexual (fidelity and nucleotide discrimination) and sexual (viral RNA recombination) RNA replication mechanisms influence ribavirin sensitivity and resistance.

### L420A revertants and pseudorevertants

Because an L420A polymerase mutation exacerbates ribavirin-induced error catastrophe coincident with defects in sexual RNA replication mechanisms (18), we predicted that ribavirin-resistant poliovirus selected from L420A parental strains would acquire polymerase mutations that facilitate sexual RNA replication mechanisms. Consistent with this prediction, L420A revertants or pseudorevertants (L420I and L420V) were selected in every lineage of virus containing an L420A mutation: 3/3 lineages of L420A parental virus, 3/3 lineages of G64^Fix^ L420A parental virus and 3/3 lineages of G64S L420A parental virus (Table 1). Importantly, L420A revertants and the L420I and L420V pseudorevertants restored efficient sexual RNA replication mechanisms coincident with ribavirin resistance (Fig. 7). These data reinforce the correlation between efficient viral RNA recombination and resistance to ribavirin-induced error catastrophe (18).

### Novel ribavirin-resistance mutations

Serial passage of poliovirus or FMDV in escalating concentrations of ribavirin leads to the selection of ribavirin resistant polymerase mutations: a G64S mutation in the poliovirus polymerase (53) and an M296I mutation in the FMDV polymerase (62, 63). By using several ribavirin-sensitive and ribavirin-resistant parental viruses, we obtained both known and unknown ribavirin resistant polymerase mutations following serial passage of poliovirus in escalating concentrations of ribavirin (Fig. 2 & Table 1). G64S mutations were selected twice, M323I was selected twice, and M299I, T353I and M392V were each selected once. The M299I mutation in poliovirus is distinct from the ribavirin-resistant M296I mutation selected in FMDV as the poliovirus M299 residue corresponds to FMDV-I309 (62, 63). Our biochemical data indicate that polymerase mutations selected during serial passage in ribavirin tend to have reduced rates of RNA elongation and increased discrimination factors (Table 3). M299I, M323I and T353I mutations reduced the rate of RNA elongation in processive elongation assays from 26 to ≈18 nt per sec, with slight increases in the fidelity of RNA synthesis (Discrimination Factors of 120 ± 10 for M299I polymerase, 130 ± 10 for M323I polymerase and 160 ± 10 for T353I polymerase).

The M392V polymerase mutation selected during serial passage in ribavirin was confirmed to mediate ribavirin-resistance in poliovirus re-derived from an infectious cDNA clone (Fig. 7A). This mutation increased the polymerase Discrimination Factor (Table 3) without changing the frequency of viral RNA recombination (Fig. 7B), consistent with an impact on the fidelity of viral RNA synthesis. Curiously, a mutation at an analogous site in the EV-71 polymerase (M393L) is reported to resist the antiviral drug NITD008 (64). NITD008 is an adenine analogue whereas ribavirin is a general purine analogue. Both NITD008 and ribavirin are pro-drugs converted into triphosphate forms in vivo where they function as NTP substrates in the active site of the polymerase. Deng and colleagues note that the EV-71 M393 side chain interacts with the primer grip of the polymerase (64). They suspect that the M393L mutation influences the polymerase active site in a way that inhibits NITD008 antiviral activity. We find that a M392V mutation decreased the rate of RNA elongation coincident with an increase in the poliovirus polymerase Discrimination Factor (Table 3) without changing the frequency of viral RNA recombination (Fig. 7B). These data suggest that the M392V mutation can influence the rate of catalysis in the active site of the polymerase, consistent with the conclusions of Deng and colleagues (64).

Two important phenotypes were shared by all of the ribavirin-resistant polymerase mutations in poliovirus: efficient viral RNA recombination (Fig. 7) and increased fidelity of RNA synthesis (Table 3). A G64S mutation renders poliovirus resistant to ribavirin-induced error catastrophe by increasing the fidelity of RNA synthesis (53, 58), but a G64S mutation is insufficient for ribavirin resistance when viral RNA recombination is disabled by an L420A mutation (18). Consequently, poliovirus with both G64S and L420A polymerase mutations is highly sensitive to inhibition by ribavirin (Fig. 7, G64S L420A). Furthermore, when subjected to serial passage in ribavirin, the L420A mutation in G64S L420A virus reverted to L420 in 3/3 lineages (Table 1), resulting in ribavirin-resistant G64S poliovirus.

Remarkably, despite substantial selective pressure, neither wt parental strains nor G64S parental strains acquired any new polymerase mutations during serial passage in escalating concentrations of ribavirin (Fig. 2 and Table 1). We used an MOI of 0.1 PFU per cell for each serial passage in ribavirin to avoid genetic bottlenecks. Under these conditions, ribavirin resistant variants only become fixed in the population if they outcompete the parental strain from which they arise. Serial passage of poliovirus at lower MOIs might increase the selective pressure of ribavirin, as decreased virus populations are more sensitive to inhibition by ribavirin (65). Because sexual RNA replication via viral RNA recombination requires two parental templates, decreased populations of virus should reduce the frequency of viral RNA recombination, effectively mimicking the effects of an L420A mutation.

### An extended primer grip in the viral polymerase

By using a codon mutation to inhibit the rapid emergence of the G64S classic mutation, we identified M392V as a novel ribavirin resistance mutation. The M392V mutation provided modest ribavirin-resistance as compared to the magnitudes of ribavirin resistance found in the G64S virus cluster (Fig. 7A, G64S, M299I, M323I & T353I). Structurally, M392 is found between L420 and the primer grip in polymerase elongation complexes and L420, M392, and K375/R376 interact directly with viral RNA product strand at the third, second and first bases from the active site, respectively. (Fig. 8). These protein-RNA interactions at the core of the polymerase facilitate both asexual and sexual RNA replication mechanisms. A L420A mutation specifically disrupts sexual RNA replication mechanisms without inhibiting asexual RNA replication mechanisms (Fig. 1) (18, 19) while K375A and R376A mutations impaired asexual RNA replication mechanisms to a sufficient degree that both mutations were unstable (Table 2). These results are consistent with the lethal effects of clustered charge to alanine substitution mutations at these locations in the polymerase (66). More conservative substitution mutations, K375R and R376K, were stably maintained in virus and neither of these mutations disabled viral RNA recombination, but they had divergent effects on ribavirin sensitivity, which the R376K mutation increased whereas the K375R mutation had little impact.

**Figure 8.**
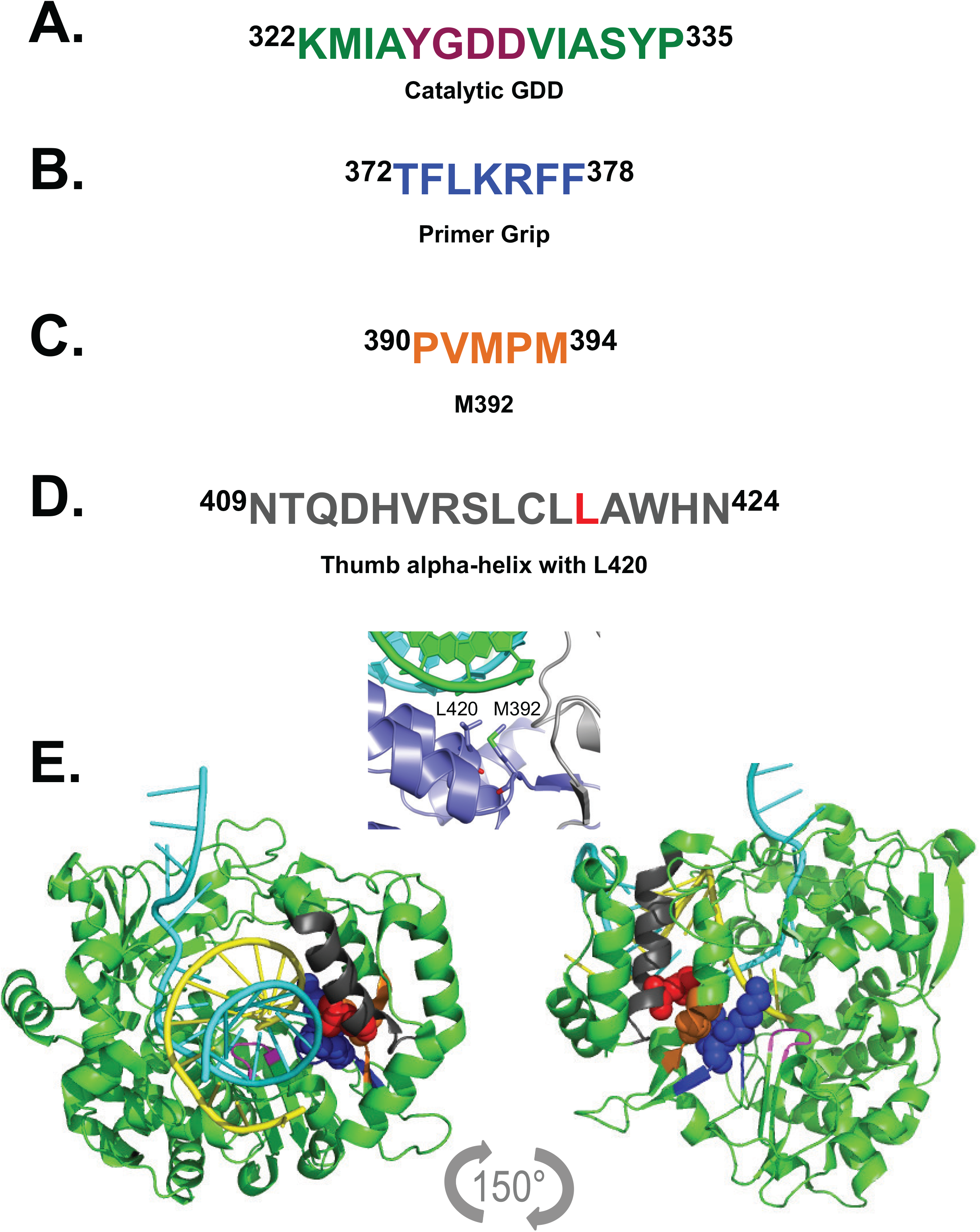
An extended primer grip in the viral polymerase mediates sexual RNA replication mechanisms. Poliovirus polymerase amino acid sequences: the catalytic GDD residues (A), primer grip (B), M392 (C) and thumb alpha-helix with L420 (D). L420 (red), M392 (orange) and primer grip residues K375 & R376 (blue) are located next to one another, adjacent to the catalytic site in the polymerase (E). L420, M392 and primer grip residues K375 & R376 constitute an extended primer grip: L420 and M392 work coordinately with the primer grip to hold nascent RNA products on homologous RNA templates near the catalytic site of the polymerase. Poliovirus polymerase PDB entry 4K4T rendered using the PyMOL molecular graphics system (Schrodinger, LLC). Polymerase is shown in green with the active site GDD residues (fuchsia), template (blue), and product (yellow) RNAs indicated in color.

Due to the interesting location of the M392V mutation within the elongation complex, we engineered two additional mutations, M392A and M392L, at this residue. Both were stably maintained in virus, but divergent phenotypic effects were observed. Virus with the M392A mutation was highly susceptible to inhibition by ribavirin, virus with the M392L mutation had no impact on ribavirin sensitivity, and virus with the M392V mutation was modestly resistant to ribavirin. None of these M392 mutations had significant impacts on the magnitudes of viral RNA recombination. However, biochemical data provided key information: the M392V mutation decreased elongation rates coincident with increased nucleotide discrimination (Table 3). These data suggest that the M392V mutation impacts asexual RNA replication mechanisms by slowing the rates of elongation coincident with increases in the fidelity of RNA synthesis.

Together, the data from the mutations at L420, M392, and the primer grip suggest that these residues form an extended primer-grip surface on the polymerase that senses the complementarity of the product duplex in the immediate vicinity of the active site (Fig. 8). Recombination involves resumption of elongation with a new primer-template pairing in the active site, and this extended primer grip would energetically favor proper Watson-Crick base pairs in dsRNA priming region, and thus facilitate recombination by selecting for proper base pairing at the critical step when elongation resumes on the new template. Consistent with this, the recombination mutations generally show reduced elongation complex stability that is indicative of impaired polymerase-RNA contacts.

### Reconciling distinct phenotypes of Y275H and L420A mutations

A L420A mutation renders poliovirus susceptible to ribavirin-induced error catastrophe coincident with defects in viral RNA recombination (18, 19). Consequently, we conclude that viral RNA recombination counteracts ribavirin-induced error catastrophe (18). In this study, we find that L420A revertants and pseudorevertants (L420I and L420V) restored efficient sexual RNA replication mechanisms coincident with increased resistance to ribavirin, as compared to L420A parental strains (Fig. 7). These data reinforce the correlation between efficient viral RNA recombination and resistance to ribavirin-induced error catastrophe (18). However, data from a Y275H mutation seem to be inconsistent with the conclusion that viral RNA recombination counteracts ribavirin-induced error catastrophe (Fig. 7, Y275H). A Y275H mutation inhibited viral RNA recombination without rendering poliovirus more susceptible to ribavirin (Fig. 7A and 7B). These results are similar to those reported by Acevedo and colleagues (60). At present, we are uncertain why the Y275H mutation inhibits viral RNA recombination without rendering poliovirus more susceptible to ribavirin. Importantly, the Y275 and L420 residues are structurally and functionally distinct. The Y275 residue is located in the fingers domain and the Y275H mutation is 4-fold slower for template binding and initiation (Table 3), suggesting a role in template binding that is mechanistically upstream of L420’s role in discriminating between homologous and non-homologous base pairing of the primer-template duplex near the active site. Thus, Y275 recruits RNA templates in a presumably sequence independent manner whereas L420 enforces sequence-dependent steps of homologous recombination. Further work will be required to better understand these distinct steps of viral RNA recombination, and their divergent effects on ribavirin sensitivity.

### Picornavirus species groups and sexual RNA replication mechanisms

The ancient origin of picornavirus RNA-dependent RNA polymerase suggests that viral RNA recombination has been a characteristic feature of viruses for billions of years (1, 3, 4). Thus, both asexual and sexual RNA replication mechanisms likely arose coordinately in ancient times. Genetic exchange is a common event among modern picornaviruses and inevitable when two related viruses with shared RNA sequence homology co-infect cells at mucosal surfaces in the respiratory or gastrointestinal tracts (41, 42), a process facilitated by bacteria (43). The extended primer grip of the poliovirus polymerase highlighted by the work presented herein is conserved across picornavirus species groups and likely mediates genetic exchange between viruses in nature. Intraspecies recombination is well documented for enterovirus species A (39, 67), species B (38, 68), species C (36) and species D viruses (69, 70), as well as rhinovirus species groups (37, 71), parechoviruses (35) and non-human enteroviruses (41). cVDPVs arise, in part, due to improved fitness from genetic exchange with non-polio group C enteroviruses in the field (46–50). Now that we appreciate the molecular and ecological conditions that contribute to genetic exchange between picornaviruses, we may be able to exploit recombination to combat human disease. Recombination-deficient oral poliovirus vaccines are one option to consider (72); however, new approaches to exploit viral RNA recombination deserve consideration.

### Summary

An extended primer grip in picornavirus polymerases contributes to the fidelity of RNA synthesis and to efficient sexual RNA replication mechanisms. This region of the polymerase interacts with nascent RNA products near the active site, effectively sensing the degree of RNA sequence complementarity between RNA products and RNA templates. With the potential to discriminate between homologous and non-homologous RNA templates, this region of the polymerase mediates sexual RNA replication mechanisms, thereby counteracting error catastrophe and contributing to genetic exchange between related picornaviruses.

## Acknowledgements

This work was supported by Public Health Service grants from the National Institutes of Health (AI059130 to O.B.P. and AI042189 to D.J.B.)

